# Development of high-affinity, single-domain protein binders for neutralizing household allergens

**DOI:** 10.1101/2025.08.03.668213

**Authors:** David KY Zhang, Qing Yue, Qiang He, Yang Li, Zihan Qu, Gary Zhao, Nathan Andreissen, Evan M Zhao

## Abstract

Feline, canine, and human dust mite allergens drive significant IgE-mediated allergies in the household environment. Here, we describe the discovery and characterization of novel protein binders, termed AVA, which efficiently bind and disrupt major allergens from cats (Fel d 1), dogs (Can f 1, Can f 2), and dust mites (Der p 1, Der p 2). Leveraging camelid single-domain variable-heavy chain (VHH) antibodies, we identified and optimized VHH sequences with robust affinity towards the individual allergens. AVA disrupted critical allergen substructures (Fel d 1) and catalytic functions (Der p 1) in molecular dynamics simulations and in *in vitro* assays, respectively. Paired with robust thermostability and non-toxic *in vitro* viability profiles, AVA represents a promising approach for downstream household allergen mitigation.

## Introduction

Allergies to common household pets and dust mites affect nearly 30% of the world’s population^1^, with greater than 90% of US households having detectable levels of multiple allergens^2,3^. Among these, cat and dog allergies are highly prevalent - roughly 70% of Americans have a pet^4^, and this number is progressively rising^5^. Due to their small size and prevalence, cat and dog allergens may remain airborne for several days with minimal air disturbance, accumulating in sofas, carpets and beds, and they may also be found in a significant (∼50%) portion of homes without pets^6^. These allergens are spread as pets groom themselves, and often settle in dust, which itself contains dust mite allergens, i.e., Der p 1 which exacerbates IgE responses when inhaled by sensitized individuals^7^.

The immune reactions towards Fel d 1, Can f 1/2, and Der p 1/2 are characterized as type I and type IV hypersensitivity, characterized by a prototypical Th2-driven IgE response produced by B cells and plasma cells. In the presence of these allergens, IgE cross-linking on sensitized mast cells and basophils occurs quickly, triggering degranulation and the release of inflammatory mediators, including histamine. The immediacy of the response correlates with the rapid onset of mild symptoms, such as itchiness of the eyes and skin, or sneezing, and more severe symptoms, ranging from perennial rhinoconjunctivitis (long-lasting nose/eye irritation), atopic dermatitis (skin irritation), asthma (difficulty breathing), and irritable bowel syndrome^1^. In rare cases, direct exposure to pets can lead to life-threatening conditions.

Of the 10 feline allergens recognized by human IgEs^1,8–12^, Fel d 1, a small ∼36 kda tetrameric glycoprotein with 2 heterodimers each comprising of two chains (chain 1: 70 amino acids; chain 2: 92 amino acids), is considered the major cat allergen that drives allergy symptoms. Fel d 1 is derived from the secretoglobin family and primarily produced in the saliva and sebaceous glands of felines^13–15^. Approximately 90% of people with cat allergies show a reaction to Fel d 1^15^-whereas the remaining allergens only show reactivity in 10-40% of allergic individuals^15,16^, with correspondingly lower levels of circulating reactive IgE. Dog allergies are driven by major dog allergens Can f 1 and Can f 2, two key members of the six described dog allergens recognized by human IgEs^17^. Can f 1 and Can f 2 are ∼27 kDa lipocalins, which feature β-barrel structures enclosing hydrophobic ligand-binding pockets that play a role in lipid metabolism^17^. 50-64% of individuals with dog allergies are sensitized to Can f 1 - among these sensitized individuals, 80% exhibit severe asthma^18^. House dust mite (HDM) allergens are ubiquitous, with at least 84% of homes testing positive^19^. Der p 1, a ∼24 kDa potent cysteine protease^20^ and Der p 2, a ∼15 kDa NPC2-family protein with structural homology to MD-2^21^ are the major dust mite allergens. These allergens are primarily localized in HDM fecal particles. In addition to mediating IgE-dependent reactions, Der p 1 disrupts epithelial tight junctions through its cysteine protease function, facilitating allergen penetration, whereas Der p 2 enhances TLR4 signaling via MyD88-dependent pathways by mimicking MD-2 and promoting LPS recognition^22^. Together, Der p 1 and Der p 2 act synergistically to amplify IgE responses in a feed-forward loop, intensifying allergic inflammation.

Allergic individuals are advised to avoid allergen exposure by removing potential allergen-containing or contaminated objects in the household. Environmental interventions, e.g., HEPA air filters, frequent and intensive cleaning, pest control, and dehumidification, are encouraged and may help alleviate symptoms^23,24^. Additionally, common over-the-counter interventions, notably antihistamines and corticosteroids, are often used. Modern approaches including allergen-specific immunotherapy, may be prescribed, but involve the risk of adverse effects and a treatment regimen requiring several dozen injections over 3 years^25^. Lower risk allergen delivery routes (e.g., sublingual, epicutaneous) and adjuvanted formulations, have also been explored and have some clinical benefit^26^. Notably, Orengo et al. developed a monoclonal IgG antibody therapy that competitively prevented Fel d 1 from binding to IgE in humans, showing good clinical benefit^27^. Thoms et al. developed a vaccine using cucumber mosaic virus-like particles (CuMV) to immunize cats, with the goal of inducing neutralizing antibodies against Fel d 1, thereby reducing its bioavailability and allergenic potential in humans^28^. These approaches can alleviate allergy symptoms in allergic individuals but often demand substantial lifestyle adjustments and possible pet relinquishment.

In the present study, we leveraged single-domain antibody fragments derived from variable heavy chain-only antibodies (VHH) to develop high-affinity protein binders to efficiently bind Fel d 1, Can f 1/2, and Der p 1/2. VHH are a unique immunological adaptation naturally found in camelids such as alpacas and llamas - they have been used in a wider range of industrial and pharmaceutical applications, including filtration of biologics from the water supply^29,30^ and as a therapeutic for the treatment of both solid tumors, including breast, lung, and ovarian cancer, and blood cancers like lymphoma and myeloma^31,32^. Thus, their biodistribution, pharmacokinetics, and pharmacodynamics in animals and humans have been extensively studied^33^. They are generally classified as non-immunogenic^34^ and offer several advantages over conventional antibodies, namely their small size (∼13kDa), high stability, and robust binding affinities^35^. We termed these engineered proteins AVA (**A**llergen-targeted **V**ariable heavy chain-only **A**ntibodies). AVA bound and disrupted their cognate allergens at nanomolar affinity. Mechanistic studies revealed that anti-Fel d 1 AVA converted 98% of the immunogenic Fel d 1 tetramer to AVA–Fel d 1 complexes monomers while anti-Der p 1 AVA reduced Der p 1 protease activity up to 93%. These findings position AVA as a proof-of-principle platform for targeted disruption of major household allergens, with a view towards practical household applications, such as sprays or filters.

## Results

### Identification and characterization of AVA

To develop AVA, we first generated full length recombinant his-tagged Fel d 1, Can f 1, Can f 2, Der p 1, and Der p 2 and immunized alpacas with the individual allergens over 8 weeks in biweekly rounds of immunization (Supplementary Fig 1a-c). Following immunization, we constructed yeast display VHH libraries from the peripheral blood mononuclear cells (PBMCs) isolated from the immunized animals. These libraries were screened against cognate recombinant allergens Fel d 1, Can f 2, Can f 2, Der p 1, and Der p 2 in successive rounds of biopanning to enrich for single clones and high-affinity, single-domain VHH candidates (Fig. 1a). The sequences of these allergens used for immunization are shown in Table 1. We observed robust allergen-specific serum VHH responses to the immunized allergens after an average of 3 rounds of immunization (Supplementary Fig. 1d-f). Animals immunized with the Der p 1 allergen required an additional two rounds of immunization before reaching a sufficient titer suitable for VHH library construction (Fig. 1b-d). The VHH libraries comprised, on average, >109 unique clones (Supplementary Fig. 1g).

**Table 1:**
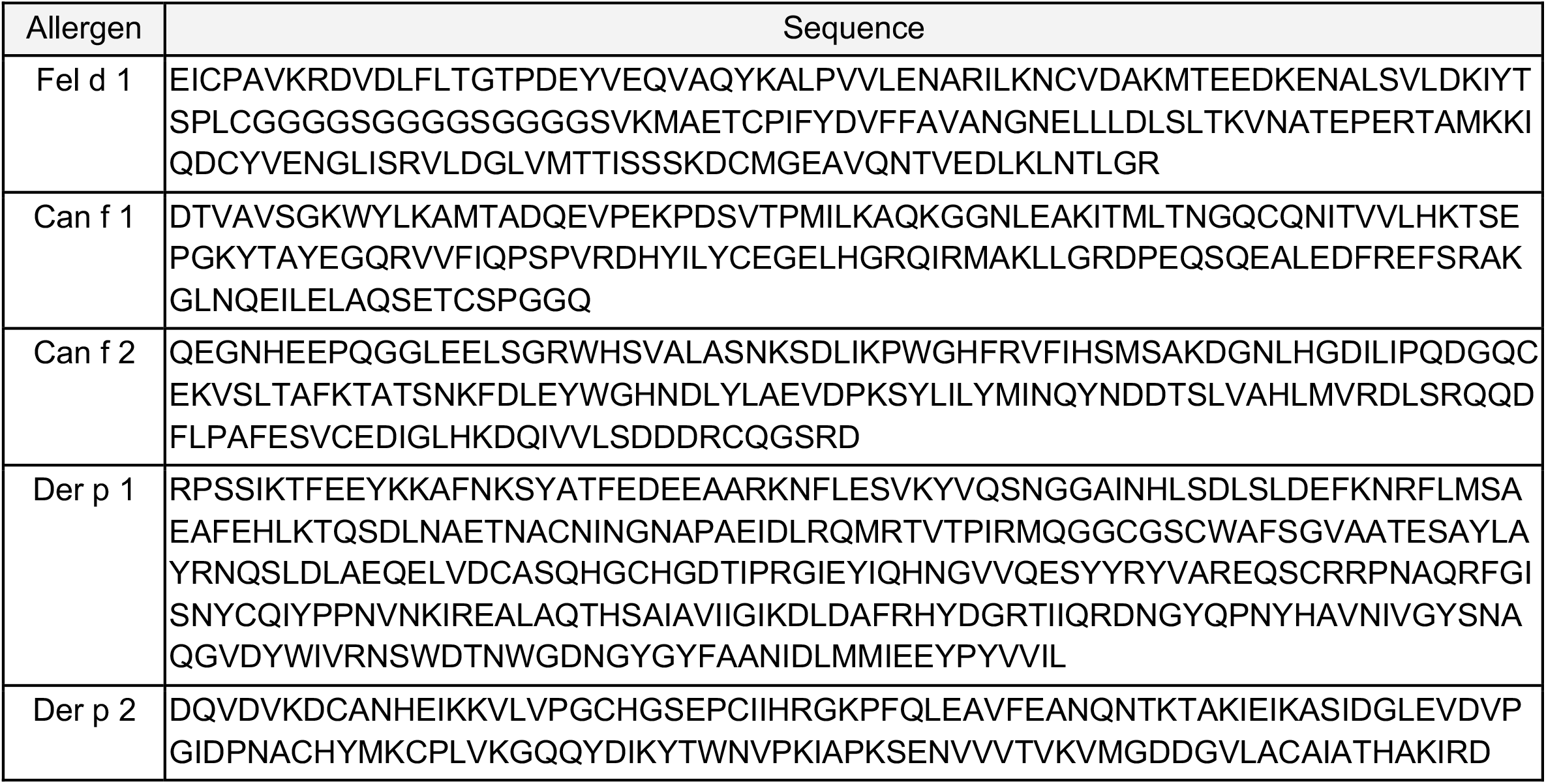
Protein sequence of allergens used in immunization studies.

**Figure 1:**
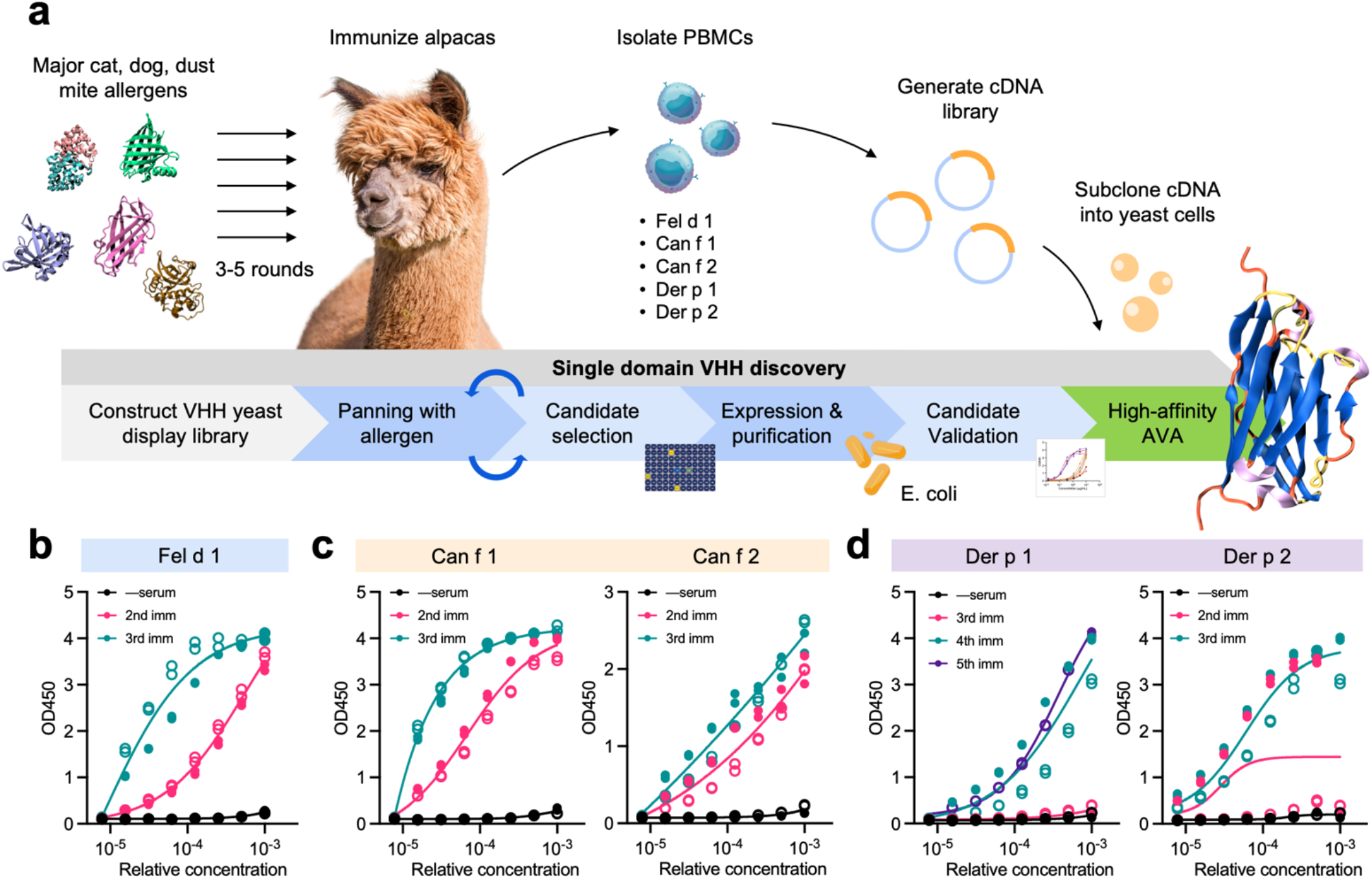
Discovery of AVA for disruption of common household allergens. (a) Schematic outline of VHH discovery, VHH candidate identification, characterization, and validation. Alpacas were immunized with 250 μg of recombinant allergen (i.e., Fel d 1, Can f 1, Can f 2, Der p 1 or Der p 2) over 3 rounds on days 0, 14, 28, and 49. For Der p 1, alpacas were immunized for an additional 5th round on day 56. For each allergen, after the last round of immunization, PBMCs were pooled from the alpacas, isolated, and their RNA was extracted and reverse-transcribed into cDNA. The VHH sequences were amplified from the cDNA and subcloned into a yeast display vector and subsequently transformed into EYB100 competent cells to construct single domain yeast display libraries. VHH candidates were identified through a series of biopanning experiments - top VHH candidates were designated AVA. (b) Serum titers of allergen-specific VHH for Fel d 1 (b), Can f 1 and Can f 2 (c), and Der p 1 and Der p 2 (d) after 2-5 rounds of immunization. Data in (b-d) represents the mean of n=2 replicates from 2 alpacas (closed/open circles).

VHH candidates specific for Fel d 1, Can f 1, Can f 2, Der p 1, and Der p 2 were selected through multiple rounds of biopanning on biotinylated target allergens by FACS (Fig. 2a). Additional rounds of biopanning were conducted for known epitopes specifically in the case of the Fel d 1 allergen. Four previously characterized and distinct epitopes from Fel d 1 were used in additional biopanning and VHH selection: Fel d 1 25-38AA, 46-59AA, and 36-49AA from chain 1, and 15-28AA in chain 2 (Supplementary Fig. 2a). Indeed, in an analysis of 61 patients with cat allergies, 65% of patients exhibited IgE binding to at least one of the peptides: 46% showed IgE binding to peptide 25-38AA and 11% showed binding to peptide 46-59AA in chain 1, while 28% showed binding to peptide 15-28AA in chain 2^36^. The VH+VL IgG sequences for Fel d 1 36-49AA have been published^37^ and were synthesized as benchmark controls (370VH+378VL, 18VH+378VL; Supplementary Fig. 2b). Novel VHH candidates identified in these separate rounds were consolidated with the original set of anti-Fel d 1 VHH candidates. Biopanning with biotinylated recombinant Fel d 1, Can f 1, Can f 2, Der p 1, or Der p 2 yielded single clones and subsequent sequencing analysis revealed differential VHH sequences for each allergen. For Fel d 1, 134 clones were sequenced, yielding 26 unique sequences. For Can f 1, 22 differential sequences were identified from 41 clones. For Can f 2, 21 differential VHH sequences were identified from 101 clones. For Der p 1, 180 unique were identified and sequenced, and 30 differential VHH sequences were identified. For Der p 2, 56 clones were identified and sequenced, yielding 24 differential VHH sequences. These resulting VHH sequences were expressed in E. coli, purified and further characterized for their binding properties by ELISA. The purified VHH candidates were approximately ∼12-13 kDa in size and comprising on average 120 amino acids. Out of 49 VHH sequences (1,176 pairwise combinations), only 17 pairs (1.45%) exhibited a sequence similarity score above the threshold of 0.7, indicating a generally low level of sequence similarity across the VHH candidates (Fig. 2b).

**Figure 2:**
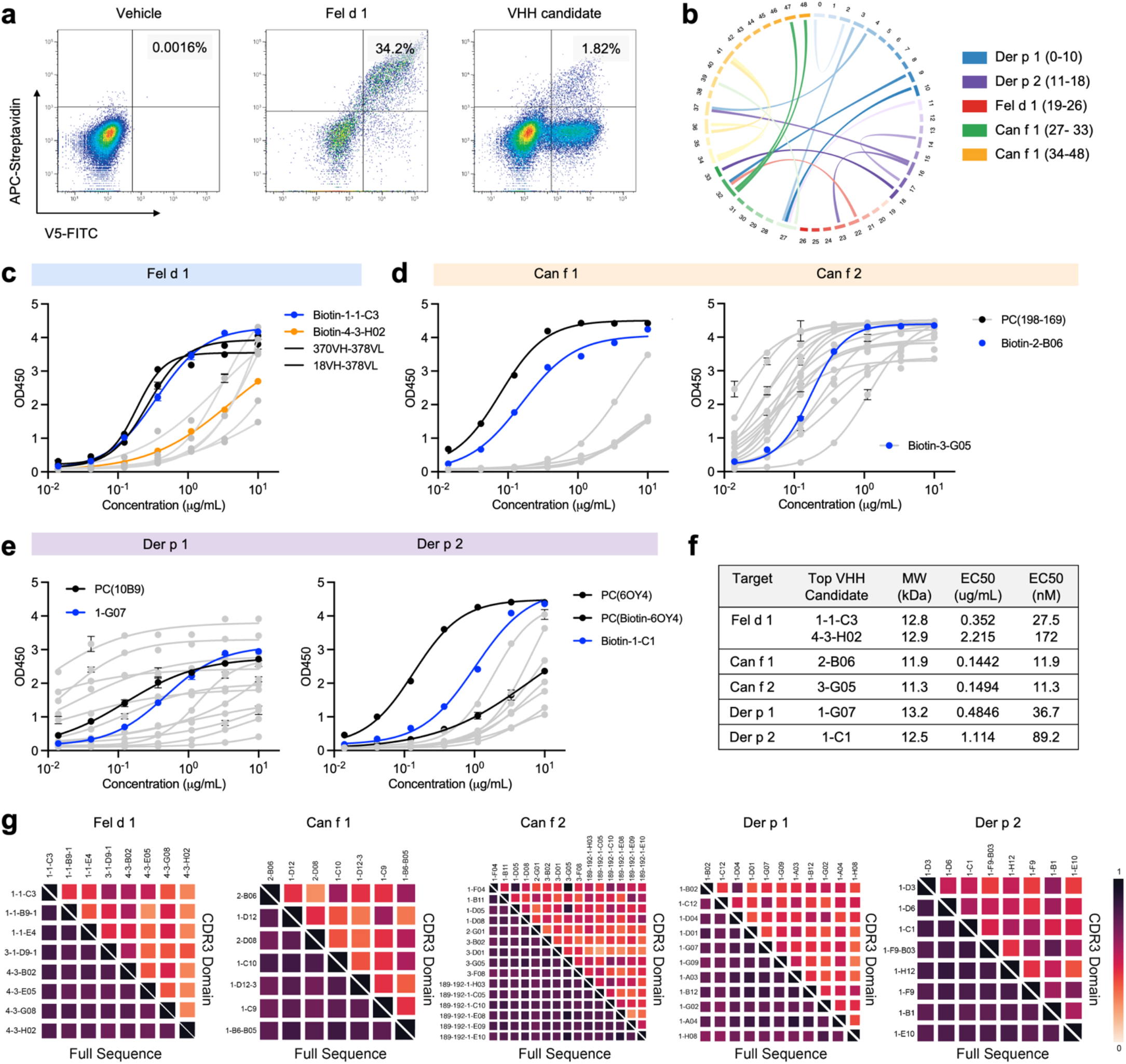
Identification of high-affinity AVA. (a) Representative FACS plots of biopanning yeast-displayed VHH with biotinylated Fel d 1. APC-streptavidin was used to visualize VHH and V5-FITC was used to validate VHH production. Left: vehicle control, middle: yeast-displayed anti-Fel d 1, right: representative VHH candidate. (b) Chord diagram of the VHH candidates’ CDR3 sequences. A total of 49 VHH candidates are shown: 8 anti-Fel d 1 VHH, 7 anti-Can f 1 VHH, 15 anti-Can f 2 VHH, 11 anti-Der p 1 VHH, and 8 anti-Der p 2 VHH. The sequence similarity was between different VHH specific for cognate allergen targets. Dose curves of VHH candidates against their cognate allergens: Fel d 1 (c), Can f 1 and Can f 2 (d), and Der p 1 and Der p 2 (e). The colored lines represent the selected top VHH candidates. (f) Summary of EC_50_ of top VHH candidates. (g) CDR3 amino acid diversity analysis of VHH candidates for each allergen. Significant variation was observed in the CDR3 region of all VHH. Data in (c-e) represents mean ± s.d. and are representative of two experimental replicates.

For each allergen, we selected 1-2 top VHH candidates based on their binding affinity towards their cognate allergen across a >3 log concentration range (Fig. 2c-e). For cat allergen Fel d 1, we identified 2 top VHH candidates (AVA-1-1-C3 and AVA-4-3-H02), which were selected from 9 VHH candidates. The anti-Fel d 1 IgGs (i.e., 370VH+378VL and 18VH+378VL) were used as benchmarks. For dog allergens Can f 1 and Can f 2, we identified a single top VHH candidate (i.e., AVA-2-B06 and AVA-3-G05) each from a selection of 7 VHH candidates (Can f 1) and 15 VHH candidates (Can f 2), respectively (Fig. 2d). We did not use a benchmark IgG for Can f 1 or Can f 2, instead opting for reactive alpaca serum due to availability. For dust mite allergens Der p 1 and Der p 2, we identified a single top VHH candidate (i.e., AVA-1-G07 and AVA-1-C1) each from a selection of 11 VHH candidates (Der p 1) and 8 VHH candidates (Der p 2), respectively (Fig. 2e). The monoclonal IgGs 10B9 (anti-Der p 1) and 6OY4 (anti-Der p 2) were used as benchmarks. The selected top VHH for each allergen consistently exhibited robust binding affinities in the nanomolar range (Fig. 2f). Comparing the VHH candidates for each allergen, high sequence diversity was observed in the CDR3 region (Fig. 2g). The anti-Der p 1 and anti-Der p 2 VHH candidates had relatively more similar CDR3 regions than anti-Fel d 1, anti-Can f 1, or anti-Can f 2 VHH. Together, these data demonstrate the discovery of high-affinity, single-domain protein binders against major cat, dog, and dust mite allergens.

### Mechanism of AVA–allergen disruption

Next, we explored the mechanism by which AVA binds and disrupts allergens, focusing on the Fel d 1 allergen, given its relatively high prevalence in the household environment^1^. Fel d 1 may be found at an average of 4.73 µg/g^6^ compared to Der p 1 at 1.40 µg/g^19^ and Can f 1 at 4.69 µg/g^6^. We conducted a series of computational investigations characterizing the interaction between AVA-1-1-C3 (a top VHH candidate) and Fel d 1 (Fig. 3a-d). The initial binding configuration was predicted using AlphaFold 3, and its stability was subsequently validated through an extended molecular dynamics (MD) simulation with annealing. Interfacial root-mean-square deviation (RMSD) analysis confirmed the stability of the protein complex. AlphaFold 3 predicted a binding configuration for the Fel d 1–AVA-1-1-C3 complexes, revealing the target Fel d 1 binding site as 36-49 AA. This prediction is consistent with the trajectory analysis, supporting our previous findings. Two AVA-1-1-C3 molecules were found to bind symmetrically at the dimer-dimer interface of the Fel d 1 tetramer, forming a stable and symmetric complex (Fig. 3a). The 37, 52-54, 59-61AA of AVA-1-1-C3 engages directly with the Fel d 1 binding site, while the loop spanning 101-107AA extends into the binding interface of the tetramer (Fig. 3b). This loop insertion disrupts the native binding architecture of the Fel d 1 tetramer. Contact map analysis revealed that upon AVA-1-1-C3 binding, 32 amino acids that previously participated in the interface were displaced, whereas 16 new residues become involved in inter-subunit interactions (Fig. 3c). Among the 32 displaced AA, AA31 to 34, 86, 89, 91, 93, 94 directly bind with AVA-1-1-C3. These results suggest that binding of AVA-1-1-C3 substantially disrupts the native configuration of the Fel d 1 tetramer, facilitating its destabilization and likely, its subsequent degradation.

**Figure 3:**
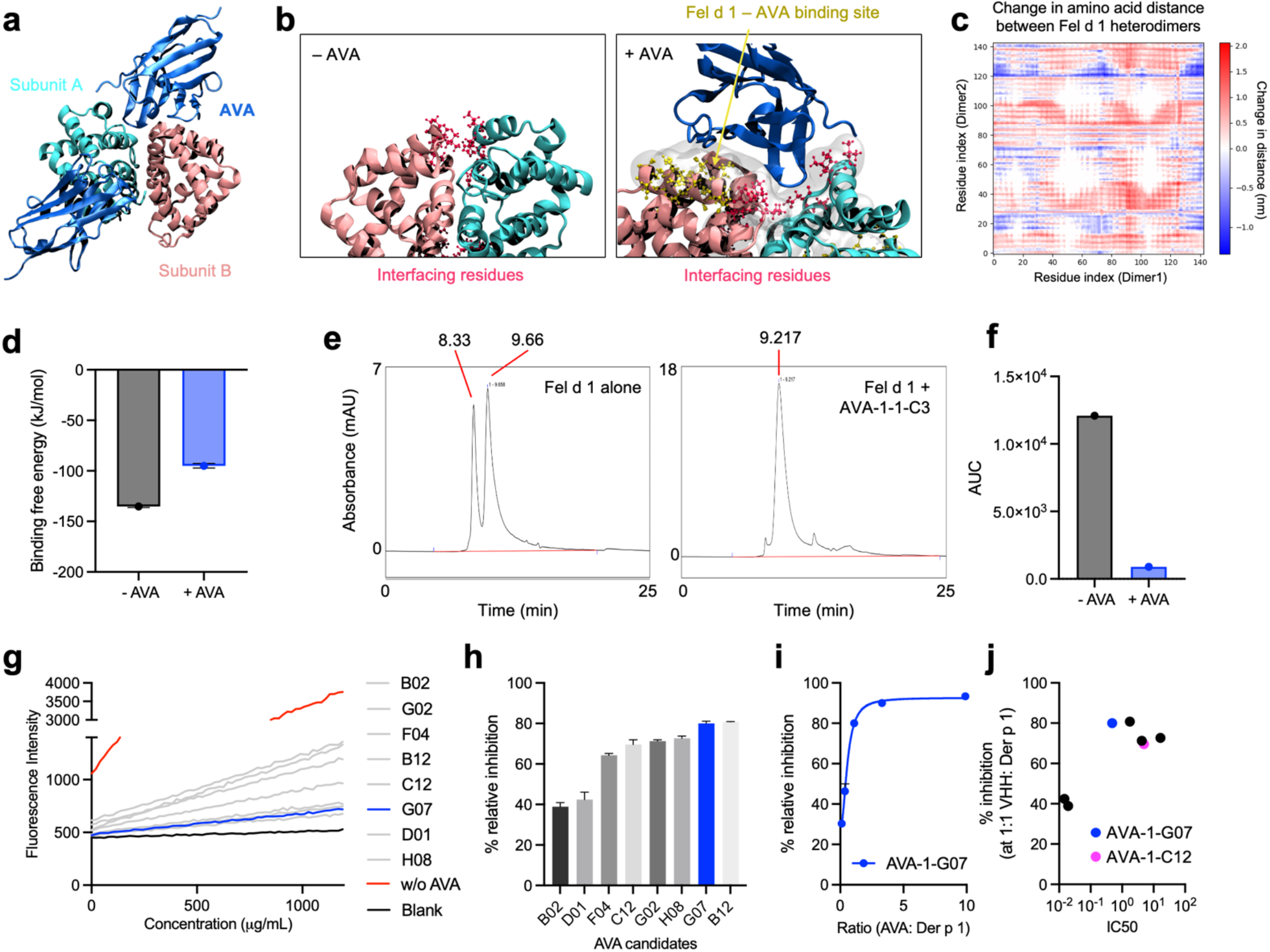
AVA binds and disrupts their cognate allergens. (a) Overall structure of the Fel d 1 tetramer in complex with AVA-1-1-C3. Fel d 1 dimers are shown in pink and cyan; AVA-1-1-C3 is shown in blue. (b) Binding interface between AVA-1-1-C3 and the Fel d 1 tetramer. AVA-1-1-C3 binds stably at the Fel d 1 binding site, with residues 101-107 extending into the tetramer interface and disrupting dimer-dimer interactions. Green: Fel d 1 binding site; yellow: residues that now bind to AVA-1-1-C3 but previously interacted with the opposing dimer; gray: the interface between Fel d 1 and AVA-1-1-C3. (c) Changes in inter-dimer contacts upon AVA-1-1-C3 binding. Red: residues that previously participated in dimer–dimer binding but no longer do so; blue: residues that now engage in inter-dimer interactions but were not involved previously. (d) Binding free energies between Fel d 1 dimers in the absence and presence of AVA-1-1-C3. (e) Size exclusion chromatograms on free Fel d 1 (e, left) and of Fel d 1 pre-incubated with AVA-1-1-C3 (e, right). Fel d 1 was primarily observed as smaller dimers (peak at 9.66) and larger tetramers (peak at 8.33) but was almost entirely converted to Fel d 1-AVA-1-1-C3 complexes (peak at 9.217) when incubated at a 1:1 molar ratio. (f) Area under the curve quantitation of tetrameric Fel d 1 and monomeric Fel d 1-AVA-1-C3 complexes. (g) Kinetic change in the fluorescence intensity of the fluorogenic substrate following treatment with AVA. (h) Relative % inhibition of Der p 1 cysteine protease activity at a 1:1 molar ratio of VHH:Der p 1. (i) Relative % inhibition at varying VHH: Der p 1 ratios for the lead VHH candidate AVA-1-G07. (k) Scatter plot comparing the binding EC_50_ against the relative % inhibition of Der p 1 at the 1:1 molar ratio of VHH:Der p 1. Data in (d, h) represents mean ± s.d.

To confirm these observations, we evaluated the binding affinity of AVA-1-1-C3 by performing a series of umbrella sampling MD simulations. We observed that the binding free energy between the Fel d 1 dimers within the tetramer is approximately −135.25 kJ/mol and umbrella sampling simulations of the AVA-1-1-C3–Fel d 1 tetramer complex revealed that the inter-dimer binding energy is reduced to −94.96 kJ/mol in the presence of AVA-1-1-C3 (Fig. 3d). These findings suggest that AVA-1-1-C3 weakens inter-dimer interactions, facilitating the disassembly of the Fel d 1 tetramer (Supplementary Fig. 3). To validate these simulations, we performed a size-exclusion chromatography (SEC) study and incubated Fel d 1 with AVA-1-1-C3 at a 1:1 mass ratio prior to measurement. We observed that Fel d 1 alone existed primarily in its tetrameric and heterodimeric forms (Fig. 3e). However, following the addition of AVA-1-1-C3, only AVA-1-1-C3–Fel d 1 heterodimers were observed, representing a 98% conversion from free Fel d 1 to AVA-1-1-C3–Fel d 1 based on the area under the curve, confirming our simulation results (Fig. 3f).

Next, we turned our attention to Der p 1. In addition to mediating IgE-dependent responses, Der p 1’s cysteine protease activity is an important contributor to its immunogenicity and has shown to augment IgE responses to itself and other allergens^7^. To validate the bioactivity of AVA-1-G07, our top anti-Der p 1 VHH candidate, we incubated the protein with Der p 1 and determined the enzyme’s change in cysteine protease activity using a continuous rate assay. We included an additional 8 anti-Der p 1 VHH candidates for comparison (Fig. 3g). We observed varying levels of Der p 1 protease activity inhibition across the VHH candidates (Fig. 3h), but a robust, dose-dependent deactivation of Der p 1 protease activity with AVA-1-G07 (Fig. 3i). At a 1:1 molar ratio of AVA-1-G07 to Der p 1, an 80% reduction in Der p 1 protease activity was observed, which increased to a maximum reduction of 93% at a 10:1 molar ratio. Interestingly, we observed a poor correlation between the anti-Der p 1 VHH’s binding EC_50_s and the % of Der p 1 cysteine protease inhibition at the 1:1 molar ratio, suggesting that some VHH candidates likely interacted with non-catalytic domains of the Der p 1 enzyme (Fig. 3j). However, we observed that AVA-1-C12, which showed the greatest relative sequence similarity to AVA-1-G07 but worse EC_50_ (Fig. 2g), strongly inhibited Der p 1 protease activity, suggesting the two VHH share the same target epitope. Overall, these data suggest that a key mechanism by which AVA-1-G07 neutralizes Der p 1 is by inhibiting its cysteine protease activity.

### In vitro profile of AVA

Next, we confirmed the thermostability and cytotoxicity profile of the AVA. We investigated the thermostability of various AVA, focusing on the top VHH candidates we previously characterized. We observed that all AVA demonstrated T_m_ > 50 ℃ and T_agg_ > 40 ℃ (Fig. 4a), surpassing typical thresholds of developable antibodies and accelerated storage conditions, respectively^38^. As an example, AVA-1-1-C3 exhibited robust integrity, with aggregation (T_agg_) observed at 58.9 ℃ and melting (T_m_) observed at 66.5 ℃. The high T_agg_ and T_m_ suggested a low propensity of AVA to aggregate or unfold under ambient conditions, indicating that AVA exhibited robust thermostability, as expected. The T_m_ and T_agg_ of other AVA are shown in Table 2. Next, we conducted a series of *in vitro* toxicity studies using AVA-1-1-C3 as a representative AVA. A selection of common primary human skin cells, specifically human dermal fibroblasts (HDF), human epidermal keratinocytes (HEK), and human follicle dermal papilla cells (HFDPCs) were treated with AVA-1-1-C3 across a >3 log concentration range. Non-skin cells, including human dermal microvascular endothelial cells (HDMEC), human skeletal muscle cells (HSKMC), and human renal cortical epithelial cells (HRCEpC) were also tested (Fig. 4b-g). Using bovine serum albumin (BSA) as a benchmark, we only observed a minor decrease in cell viability for both BSA and AVA-1-1-C3 at very high concentrations (78 µM), which was likely due to the substantial alterations in culture media. In human skin dermal follicle papilla cells (HFDPC; Fig 4d), the decrease in cell viability was more pronounced, but was present in both AVA-1-1-C3 and BSA groups, suggesting that HFDPC may be more innately sensitive to changes in the culture media. In non-skin cells, HSKMC, HRCEpC, and HDMEC, no substantial change in cell viability was observed across the same >3 log concentration range of AVA-1-1-C3 or BSA. These data highlight that AVA are non-cytotoxic to primary human cells.

**Table 2:** Thermostability of AVA.

| Allergen target | AVA protein | T<sub>agg</sub> (°C) | T<sub>m</sub> (°C) |
| --- | --- | --- | --- |
| Fel d 1 | 1-1-C3 | 58.9 | 66.5 |
| Can f 1 | 2-B06 | 48.0 | 56.4-63.7 |
| Der p 2 | 1-C1 | 43.5 | 54.5-66.5 |

**Figure 4:**
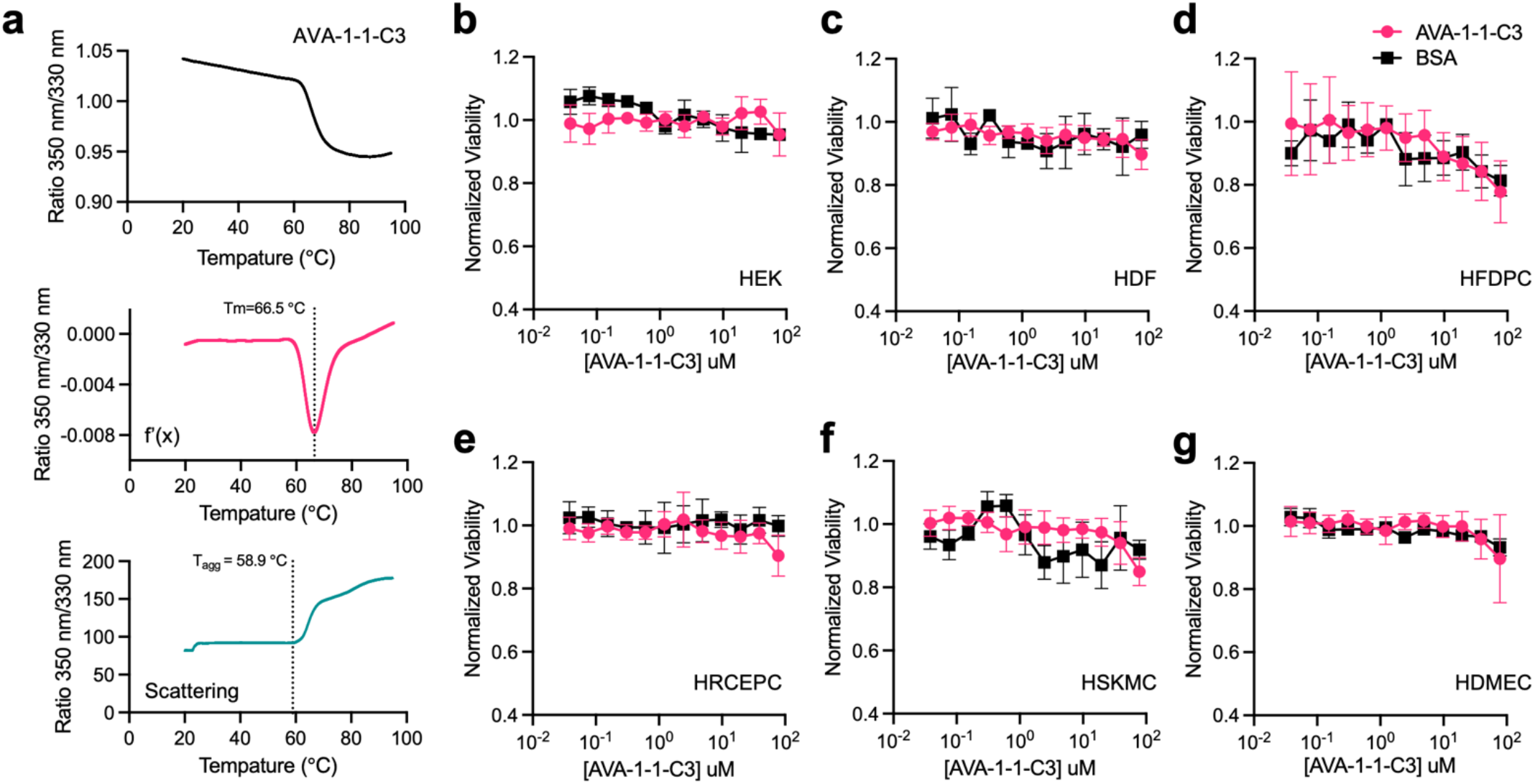
AVA are temperature stable and non-toxic. (a) Thermostability profile of AVA-1-1-C3 as a representative protein, as determined by differential scanning fluorimetry. (b-g) Cytotoxicity of primary human cells following 48-hour exposure to AVA-1-1-C3 or an equivalent concentration of bovine serum albumin (BSA), evaluated using the CellTiter-Glo® (CTG) luminescent cell viability assay. Cell types included human epithelial keratinocytes (HEK, b), human dermal fibroblasts (HDF, c), human follicle dermal papilla cells (HFDPC, d), human renal cortical epithelial cells (HRCEPC, e), human skeletal muscle cells (HSKMC, f), and human dermal microvascular endothelial cells (HDMEC, g). No significant differences in cell viability were observed. Data in (b-g) represents mean ± s.d. and are representative of two experimental replicates.

## Discussion

Our study demonstrates that engineered VHH (termed AVA) can robustly bind and disrupt major household allergens from cat, dog, and house dust mites, offering a novel tool to manage environmental allergens. Current approaches to environmental allergen control, including HEPA filtration or rigorous cleaning regimens, have struggled to consistently reduce allergen burden or sufficiently alleviate allergy symptoms for sensitized individuals^24,39^. The utility of allergen-specific immunotherapies, while curative for many, are constrained by lengthy treatment regimens, risk of adverse reactions, and variable efficacy^40^. Monoclonal anti-Fel d 1 IgG^27^ can block IgE binding but do not alter Fel d 1 allergen structure, and dietary IgY supplementation for felines may limit allergen shedding but offers limited control over allergens already dispersed throughout the home^41^.

AVA acts at the critical intersection of high-affinity allergen binding and direct structural and functional disruptions: AVA-1-1-C3 destabilizes two Fel d 1 heterodimers; computational modeling and biophysical assays confirmed that AVA-1-1-C3 binds at Fel d 1’s dimer-dimer interface, causing a robust 40.3 kJ/mol change in tetramer binding affinities. For context, the binding affinities between HER2 and trastuzumab is −46.0 kJ/mol, and −47.4 kJ/mol for rituximab binding to CD20^42,43^. It is possible that AVA–Fel d 1 complexes could retain residual immunogenicity; this will be addressed in future immunogenicity studies. AVA-1-G07 inhibits the cysteine protease activity of Der p 1 - two mechanisms directly linked to allergenicity and IgE-sensitization potential. For Der p 1, the lack of correlation between binding affinity and protease inhibition among VHH candidates suggests that epitope specificity, i.e., targeting Der p 1’s catalytic domain, may be critical for neutralization. AVA are ideal candidates for potential downstream application: their thermostability, low aggregation propensity, and their non-toxicity to diverse skin and non-skin primary human cell types suggest a strong capacity for the translation of molecular mechanisms to downstream user benefit.

However, several caveats and limitations warrant discussion. While *in vitro* and computational data support AVA-mediated disruption of cat, dog, and dust mite allergens, these observations require further validation in human studies. Moreover, since sensitization profiles across individuals are naturally heterogeneous, expanding the AVA repertoire to include additional allergens such as Fel d 4 (one third of allergic individuals with cat allergies are sensitized to Fel d 4^16^) or Can f 5 may enhance population coverage. Additionally, little is known about the specific biological functions for Fel d 1/2 and Can f 1/2, rendering *in vitro* functional characterization challenging. Our findings highlight AVA as a proof-of-principle modality for binding and disruption of major household allergens. Future studies will focus on translating these findings into practical applications. By shifting the focus from symptomatic treatment and allergen avoidance to targeted neutralization of allergen triggers in the home, such an approach has the potential to benefit millions of Americans affected by pet and dust mite sensitization.

## Materials and Methods

### Reagents and materials

Reagents for enzyme-linked immunosorbent assay (ELISA) included HRP-conjugated rabbit anti-Llama IgG (H+L) secondary antibody (Novus, CAT# NBP1-7509), HRP-conjugated rabbit anti-camelid VHH cocktail (Genscript, CAT# A02016), HRP-streptavidin (Boster, CAT# BA1088), HRP-protein A (Boster, CAT# BA1080), FITC conjugated V5 Tag Antibody (iCareab, cat# IAB012B, streptavidin-PE (eBioscience, CAT# 12-4317-87), streptavidin-APC (BioLegend, CAT# 405207), and streptavidin-coated magnetic beads (NEB, CAT# S1420S). For signal amplification and enzymatic assays, NHS-biotin (APExBIO, CAT# A8002), Boc-Gln-Ala-Arg-AMC · HCl (Absin, CAT# abs45133013), and MPBS (PBS with 5% non-fat milk). For molecular and cellular assays, RNAiso Plus (Takara, CAT# 9109), PrimeScript^™^ II 1st Strand cDNA Synthesis Kit (Takara, CAT# 6210B), NuHi Power Mix (NuHigh Biotechnology, CAT# NH9303), and SfiI restriction enzyme (NEB, CAT# R0123L) were used in gene expression and cloning protocols described below. The gel extraction kit (Qiagen, CAT# 28706), plasmid maxiprep kit (Tiangen, CAT# DP117), and 3M sodium acetate, pH 5.2–6.0 (Sigma, CAT# 126-96-5) were used for nucleic acid purification. For transfection and display applications, we used pYDisplay yeast display vectors (iCarEab), LVtransm transfection reagent (iCarEab, CAT# LVTran100), and OPM-293 CD05 medium (OPM Bioscience, CAT# 81075-001). Adjuvant was purchased from GERBU (CAT# 3030) for alpaca immunization studies. ColorMixed protein marker (Solarbio, CAT# RP1930) was used for SDS-PAGE protein analysis. CellTiter-Glo® 2.0 (Promega, CAT# G7573) was used for cell viability studies.

Cells used in the identification and expression of AVA proteins included Origami B (Beijing Biomed Gene Technology, cat# BC219-01), 293-F (ThermoFisher, cat#A14527), and EBY100 yeast cells (Weidibio, cat# YEL1110M). For *in vitro* toxicity studies, primary cells including human dermal fibroblasts (HDF, Cell Applications, Cat# 106K-05a, lot 1632), human epidermal keratinocytes (HEK, Cell Applications, Cat# 102-05a, lot 2146), human dermal microvascular endothelial cells (HDMEC, PromoCell, Cat# C-12210, lot 483Z001.3), human skeletal muscle cells (HSKMC, Cell Applications, Cat# 150K-05a, lot 3507), and human renal cortical epithelial cells (HRCEpC, Promocell, Cat# C-12660, lot 501Z019.23). Cells were cultured in their corresponding complete media including HDF growth medium (Cell Applications, Cat# 116-500), human EpiVita serum-free growth medium (Cell Applications, Cat# 141-500a), microvascular endothelial cell growth medium (PromoCells, Cat# C-22120), HSkMC growth medium (Cell Applications, Cat# 151-500), and renal epithelial cell growth media (PromoCell, Cat# C-26130). General laboratory consumables included 50 mL Falcon tubes (Corning, Cat# 352070), 15 mL Falcon tubes (Corning, Cat# 430052), T125 shake flasks (Corning, Cat# 431143), 6-well plates (Corning, Cat# 3516), 96-well plates (Corning, Cat# 3365), RNase-free 1.5 mL Eppendorf tubes (QSP, Cat# 509-GRD-Q), RNase-free 200 μL PCR tubes (Axygen, Cat# PCR-02D-C), electroporation cuvettes (0.2 cm, Bio-Rad), and K2EDTA vacutainer blood collection tubes (BD, Cat# 367525).

### Instrumentation

A thermostatic incubator (Shanghai Jinghong, DNP-9052), CO_2_ orbital shaker (Crystal Technologies & Industries, CO-06U), CO_2_ incubator (Thermo, 3111), and clean bench (Suzhou Airtech, SW-CJ-1FD) were used for cell culture. Biosafety cabinets (Haier, HR40-IIA2) were used for cell culture studies and aseptic operations. For molecular and cellular analysis, we used a thermal cycler (Applied Biosystems, ABI2720), electroporator (Bio-Rad, 165-2661), low-speed centrifuge (Cencelab, L535R), microplate reader (Hangzhou Allsheng, AMR-100 or Promega, GloMax®), flow cytometer (Thermo Fisher, Attune Nxt), and a cell sorter (BD Biosciences, FACSAria^™^ III).

### Identification of high-affinity AVA

#### Allergen preparation

Based on the amino acid sequence of Fel d 1, Can f 1, Can f 2, Der p 1, and Der p 2 proteins (Table 1), eukaryotic expression vectors were constructed as previously described with a signal peptide (MGWSCIILFLVATATGVHS) for each sequence below^44^. After transient transfection into mammalian cells, the culture supernatant was collected, and the target recombinant protein was purified using affinity chromatography. The purity of the recombinant protein was assessed by SDS-PAGE and quality control was performed using in-house produced antibodies based on published patents (Fel d 1 : WO2013166236A1 (Heavy chain: SEQ ID NO.: 370\SEQ ID NO.: 18; Common Light Chain: SEQ ID NO.: 378);).

#### Alpaca immunization

The purified recombinant allergens Fel d 1, Can f 1, Can f 2, Der p 1, and Der p 2 were used to immunize alpacas (Lama pacos) in 14-day intervals. Beginning from the second immunization at day 28, peripheral blood was collected every seven days after each immunization to monitor the serum antibody titer. For each allergen, two alpacas were immunized and upon completion of the immunization schedule, 100 mL of peripheral blood was collected, and peripheral blood mononuclear cells (PBMCs) were isolated for the construction of a single-domain VHH display library.

#### Alpaca serum antibody titer quantification

To evaluate the immune response raised towards the immunized allergens, 5 mL of peripheral blood was collected from each alpaca and incubated at 37°C for 1 hour, followed by overnight storage at 4°C. The samples were then centrifuged at 1500 g for 20 minutes to separate the serum, which was transferred to new sterile tubes. For ELISA, the target recombinant protein (either Fel d 1, Can f 1, Can f 2, Der p 1, or Der p 2) was diluted to a final concentration of 1 µg/mL in sterile carbonate buffer saline (CBS), and coated overnight at 4°C in 96-well ELISA plates (100µL).

Following coating, the solution was removed, and plates were washed five times with PBST (0.05% Tween-20 in PBS). Each well was blocked with 200 μL of 3% non-fat milk diluted in PBS (MPBS) at 37°C for 2 hours. After removing the blocking buffer and washing five times with PBST, 100 μL serially diluted serum was added to each well and incubated at room temperature for 1 hour; PBS was used as a negative control. After incubation, the plates were washed five more times with PBST, and 100 μL HRP-conjugated anti-llama IgG (H+L) antibody (diluted 1:50,000) was added and incubated for 1 hour at room temperature. Following five more washes with PBST, 100 μL of TMB substrate was added to each well for color development in the dark at room temperature for 10–15 minutes. The reaction was stopped by adding 50 µL of stop solution per well, and the absorbance at 450 nm was measured using a microplate reader (Hangzhou Allsheng, AMR-100).

### PBMC isolation and VHH sequence cloning

#### cDNA Synthesis

A total of 100 mL of peripheral blood was collected, and PBMCs were isolated using a lymphocyte separation solution. cDNA was synthesized through reverse transcription of extracted RNA using the PrimeScript^™^ II 1st Strand cDNA Synthesis Kit. The resulting cDNA samples were either used immediately for the following experiments or stored at −20°C for later use.

#### VHH Fragment Amplification

The DNA sequence spanning the region between the signal peptide and CH2 domain of the VHH cDNA was amplified by PCR and purified via gel extraction of the ∼750 bp band. A second PCR was performed using a forward primer targeting the FR1 region and a reverse primer targeting the anti-hinge and FR4 regions containing an SfiI restriction site.

Forward primer 5’GTCCTGGCTGCTCTTCTACAAGG 3’

Reverse primer 5’ GGTACGTGCTGTTGAACTGTTCC 3’

The resulting ∼400 bp product was gel-purified. The purified PCR products were divided into 200 μL aliquots in 1.5-mL microcentrifuge tubes. To each aliquot, 20 μL of a solution containing 3 M sodium acetate and 1 μg/μL glycogen was added and mixed thoroughly by pipetting. Subsequently, 880 μL of 200-proof ethanol was added, the mixture was gently inverted to mix, and the samples were stored at −80 °C.

Forward primer 5’AGTGCAGCTCGTGGAGTCNGGNGG 3’

Reverse primer 5’ GATCACTAGTGGGGTCTTCGCTGTGGTGCG3’

### Construction of the VHH yeast display library

#### Linearization of the yeast display vector

The pYDisplay vector was linearized by the restriction enzyme SfiI at 50 ℃ overnight. The reaction mixture was resolved on 1% agarose gel followed by extraction of the band at ∼5,000 bp using the gel extraction kit. The product concentration was determined by measuring the absorbance at 260 nm using a NanoDrop spectrometer. Similar to the previous, purified linearization products were aliquoted into 200 μL and transferred to 1.5-mL microcentrifuge tubes. To each aliquot, 20 μL of a solution containing 3 M sodium acetate and 1 μg/μL glycogen was added and mixed thoroughly by pipetting. Subsequently, 880 μL of 200-proof ethanol was added, the mixture was gently inverted to mix, and the samples were stored at −80 °C.

#### VHH library construction

Yeast competent cells stored at −80 °C were streaked onto yeast extract peptone dextrose (YPD) solid medium plates and incubated at 30 °C for 3–5 days for activation. A single colony was then inoculated into 50 mL of YPD liquid medium and cultured at 30 °C with shaking at 250 rpm for 1–2 days. The linearized vector fragment and PCR product were mixed with competent cells and transferred into an electroporation cuvette for transformation. After electroporation, the yeast cells were transferred into a flask and incubated at 30 °C with shaking at 220 rpm for 1 hour to recover. Following recovery, 20 μL of the suspension was diluted 1:5,000 in synthetic dextrose plus casein amino acids (SDCAA) medium, and 100 μL of the diluted sample was plated onto SDCAA agar plates and incubated for 2–3 days to determine library titer. The remaining culture was expanded for an additional 24 hours. For long-term storage, the residual yeast culture was collected into a 50 mL centrifuge tube, centrifuged at 3,000 × g for 5 minutes, and the supernatant was discarded. The pellet was resuspended in 10 mL of SDCAA, mixed with an equal volume of 50% glycerol (1:1 SDCAA: glycerol), and stored at −80 °C and used as needed.

#### Screening of the VHH yeast display library

Yeast cells initially cultured in SDCAA medium were transferred into 250 mL shake flasks containing 50 mL of synthetic galactose plus casein amino acids (SGCAA) medium and incubated at 30 °C with shaking at 240 rpm for 16 hours. Following incubation, yeast cells were harvested by centrifugation, and the supernatant was discarded. The resulting pellets were resuspended in 1 mL of PBSA (PBS with 0.5% bovine serum albumin), transferred to 1.5 mL microcentrifuge tubes, and centrifuged at 3,000 × g for 5 minutes. After removing the supernatant, the cells were washed once more with PBSA.

In parallel, streptavidin-coated magnetic beads pre-incubated with biotinylated antigen were washed twice with PBSA (each wash conducted at 4 °C with rotation for 5 minutes) and collected using a magnetic stand. The yeast suspension was then added to the allergen-bound (Fel d 1, Can f 1, Can f 2, Der p 1, or Der p 2) beads and incubated at 4 °C with rotation for 60 minutes. After binding, the beads were collected with a magnetic stand for 15 minutes, and the unbound yeast suspension was discarded. The beads were washed three times with 0.5% PBSA under the same conditions. The washed beads were resuspended in 1 mL of SDCAA medium, and 0.5–5 μL of the suspension was diluted into 100 μL of SDCAA and plated to assess screening output. The remaining suspension was divided into two portions: one was mixed with 500 μL of 50% glycerol and stored at −80 °C until needed, while the other was inoculated into 2 mL of SDCAA medium and incubated at 30 °C with shaking at 240 rpm for 16 hours. The resulting culture was expanded into 50 mL of SDCAA in a 250 mL shake flask and incubated overnight under the same conditions. The optical density at 600 nm (OD600) was measured, and the culture was diluted in SGCAA medium to an OD600 of 1 for further overnight incubation at 30 °C with shaking. The remaining culture was mixed 1:1 with 50% glycerol and stored at −80 °C until needed.

#### FACS analysis of single clone VHH candidates

Following screening, the yeast output was plated onto SDCAA agar plates. Individual colonies were picked and cultured in SGCAA media and induced for 48 hours at 30°C with shaking at 220rpm. Induced monoclonal yeast cells were transferred to a 96-well V-bottom plate and centrifuged at 3,000 × g for 3 minutes to remove the culture medium. The cells were then resuspended in 200 μL of PBS per well and centrifuged again under the same conditions, and the supernatant was discarded. This washing step was repeated 2–3 times. Biotinylated target protein was diluted to a final concentration of 5 μg/mL, and 50 μL was added to each well to resuspend the washed yeast cells. The plate was placed on a rotator and incubated at room temperature for 30 minutes. After incubation, the cells were centrifuged at 3,000 × g for 3 minutes at room temperature, and the supernatant containing unbound protein was discarded. The cells were washed three times with PBS. Subsequently, 50 μL of APC-conjugated streptavidin (diluted 1:1,000) was added to each well, mixed thoroughly, and incubated at room temperature in the dark for 30 minutes. Following incubation, the cells were centrifuged at 3,000 × g for 3 minutes, the supernatant was removed, and the cells were washed three more times with PBS. The cells were re-suspended in 200 μL of cold PBS. Flow cytometry analysis was performed to identify positive clones binding to the target antigen. Positive clones were lysed by boiling the samples in deionized water, and the lysates were clarified by centrifugation. A 0.5 μL aliquot of the supernatant was used for sequencing, while the remaining lysate was stored at −20 °C for future use.

#### Characterization of AVA

##### Construction of prokaryotic VHH expression vector and expression and purification of VHH candidates

Single-domain VHH sequences from positive yeast clones were amplified by PCR and subsequently linearized by digestion with the restriction enzyme SfiI. The resulting fragments were ligated into the pET-21a vector to construct expression plasmids for VHH production in *E. coli* Origami B (DE3) cells. To transform the cells, a tube of competent cells was thawed on ice. Plasmid DNA was added to the cells (200 ng of plasmid for 50 μL of competent cells) followed by incubation on ice for 30 minutes with gentle mixing once or twice during the incubation. The cells were then heat-shocked at 42 °C for 75 seconds and immediately returned on ice for 2 minutes. Afterward, 500 μL of LB medium was added, and the mixture was incubated at 37 °C with shaking at 220 rpm for 30 minutes. The cells were centrifuged at 5,000 rpm for 5 minutes at room temperature, and approximately 500 μL of the supernatant was removed. The remaining 100 μL was used to resuspend the pellet, which was then spread onto pre-warmed solid agar plates and incubated overnight at 37 °C.

A single colony from freshly transformed DE3 cells, or from a glycerol stock of a monoclonal transformed cell, was inoculated into 20 mL of LB medium containing 50 μg/mL ampicillin and cultured overnight at 37 °C with shaking at ∼200 rpm. The overnight culture was diluted 1:100 into 1 L of Terrific Broth (TB) medium containing 50 μg/mL ampicillin and grown at 37 °C with shaking at 200 rpm until the OD_600_ reached approximately 0.6. Protein expression was induced with 0.2 mM IPTG, and the culture was incubated and shaken at 20 °C for 16 hours to enable VHH expression. The culture was collected and centrifuged at 5,000× g for 10 minutes at 4 °C using a high-speed refrigerated centrifuge. The supernatants were discarded, and the pellets were resuspended in 50 mM NaH2PO4, 300 mM NaCl, 10 mM imidazole, pH 7.4. The resuspended cells were lysed by high-pressure homogenizer (ATS, cat. AH-1500). The mixture was centrifuged at 5,000× g for 10 min.The supernatant was collected and filtered through a 0.45 μm membrane. A Ni-NTA resin column (5 mL column volume) was prepared and connected to a peristaltic pump and a UV detector for nucleic acid/protein monitoring. The columns were pre-washed with 5–10 column volumes of deionized water to remove storage buffer, followed by equilibration with 5–10 column volumes of PBS. The filtered sample was loaded onto the column at a constant flow rate of 1.5 mL/min using the peristaltic pump. The column was washed with wash buffer (50 mM phosphate buffer, 300 mM NaCl, 40 mM imidazole, pH 8.0) until the UV signal stabilized. Elution was performed with elution buffer (50 mM phosphate buffer, 300 mM NaCl, 250 mM imidazole, pH 8.0), and fractions were collected when an increase in UV absorbance was observed and until the signal plateaued. After elution, the column was regenerated with 5–10 column volumes of PBS. The eluted protein was dialyzed against PBS using 100-fold the sample volume. Dialysis was carried out initially at room temperature for 2 hours, followed by a buffer exchange and continued dialysis at 2-8 °C for 14-16 hours. The dialyzed proteins were collected, and the concentration was determined. If the concentration was below 0.2 mg/mL, the sample was concentrated using centrifugal ultrafiltration devices. Finally, the protein solution was filtered through a 0.22 μm sterile membrane, aliquoted into final product tubes, and stored at −80 °C.

##### Transient transfection and expression and purification of antigens

PEI transfection reagent (1 mg/mL) and the eukaryotic protein expression plasmid were thawed at room temperature and mixed thoroughly by pipetting. To prepare the transfection mixture, 2 mL of OPM-293 medium was added to one well of a 6-well plate, followed by 100 μg of expression plasmid, which was mixed thoroughly by pipetting. Then, 300 μL of PEI was added, and the mixture was immediately mixed again by pipetting and incubated at room temperature for 10 minutes. The resulting DNA/PEI complex was added to 100 mL of 293F cells at a density of 1 × 10^⁶^ cells/mL and gently mixed. The culture was incubated at 37 °C with 5% CO_2_ and agitated at 120 rpm. After five days of continuous culture, the cell suspension was centrifuged to collect the supernatant, which was filtered through a 0.45 μm membrane and transferred to a sterile centrifuge tube. The recombinant antigens were purified using nickel affinity chromatography and subsequently concentrated and buffer-exchanged using centrifugal ultrafiltration. The purified protein concentration was determined using a NanoDrop spectrophotometer, and the sample was analyzed by reducing SDS-PAGE.

##### Determining the binding affinity of VHH candidates to target allergens

The target antigen (Fel d 1, Can f 2, or Der p 1) was diluted to a final concentration of 2 µg/mL using sterile CBS. 96-well ELISA plates were coated with 100 µL per well of the diluted protein and incubated overnight at 4 °C. The coating solution was removed, and the plates were washed five times with PBST. Each well was blocked with 200 µL of 3% MPBS at 37 °C for 2 hours. After discarding the blocking buffer, the plate was washed again five times with PBST. Biotinylated recombinant antibodies were added to the wells at an initial concentration of 10 µg/mL, followed by seven 5-fold serial dilutions (100 µL per well), and incubated at room temperature for 1 hour. PBS was used as a negative control. The wells were washed five times with PBST before adding 100 µL per well of MonoRab rabbit anti-camelid VHH cocktail HRP conjugate (diluted 1:10,000), followed by another 1-hour incubation at room temperature. After washing the plate five times with PBST, 100 µL of TMB substrate solution was added to each well and incubated in the dark at room temperature for 10–15 minutes. The reaction was terminated by adding 50 µL of stop solution per well, and the OD_450_ was measured using a microplate reader.

### Computational Modeling

#### Sequence Analysis and Diversity Assessment

To evaluate diversity, pairwise alignments were first performed for all AVA protein sequences to obtain alignment scores as previously described^45^. Each score was defined such that a match between identical amino acids or nucleotides contributed +1, while mismatches contributed 0. Pairwise similarity was then calculated as:

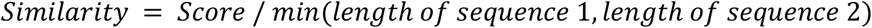

Based on the resulting pairwise similarity matrix, a chord diagram was generated using a similarity cutoff of 0.5 to visualize the diversity relationships among sequences. Subsequently, the AVA strains were categorized according to their target specificities for further analysis. For each category, sequence similarities were computed for both the full-length sequences and the CDR3 regions using the same alignment procedure described above. The CDR3 domains were extracted from all sequences using IMGT/DomainGapAlign^46^. With the similarity matrix, the similarity heatmaps were plotted.

#### AlphaFold 3 prediction of AVA-1-1-C3 structure and binding with Fel d 1 tetramer

AlphaFold 3 was used to generate the initial binding configurations of AVA-1-1-C3 with Fel d 1 tetramer. In the system, 2 entries of Fel d 1 dimer (1+2) sequences and 2 entries of AVA-1-1-C3 sequences were added. The model seeds were set to 1. The resulting structure with highest confidence was selected as the initial binding configuration. RMSD calculation was performed on the Fel d 1 tetramer to ensure a correct folding.

#### Modeling the interaction between AVA and Fel d 1

All all-atom molecular dynamics (MD) simulations were performed using GROMACS 2023.3 with the CHARMM36 force field. The folded structure of AVA-1-1-C3 was predicted using AlphaFold 3 (AF3), and MD simulations were subsequently conducted to evaluate the structural stability of the predicted model. Energy minimization, NVT and NPT equilibration, and production runs were performed while structural deviations over time were assessed using root-mean-square deviation (RMSD) analysis. The initial complex structure of AVA-1-1-C3 bound to the Fel d 1 tetramer was generated using AF3. A long production simulation with simulated annealing was applied to avoid trapping the system in a local energy minima. Energy analysis of the system showed that the total potential energy curve converged, indicating that the system had reached equilibrium and the binding between AVA-1-1-C3 and Fel d 1 tetramer is stable. The stability of the binding interface was further assessed by analyzing the root-mean-square fluctuation (RMSF) of the interface residues. Using the equilibrated structure, umbrella sampling MD simulations were conducted to estimate the binding free energy. Three systems were compared: a control system containing only the Fel d 1 tetramer, an experimental system consisting of the Fel d 1 tetramer–AVA-1-1-C3 complex, and an experimental system consisting of Fel d 1 dimer–AVA-1-1-C3 complex. The potential of mean force (PMF) profiles and corresponding binding free energies were calculated using the Weighted Histogram Analysis Method (WHAM).

#### Unbiased All-Atom Molecular Dynamics Simulations of Fel d 1 and AVA-1-1-C3

All molecular dynamics (MD) simulations were performed with Gromacs 2023.3 and CHARMM36 forcefield. Both are specially designed for simulation of biological systems. MD simulations were performed on the predicted Fel d 1 tetramer & double AVA-1-1-C3 structure. The protein was solvated in a XX x XX x XX nm^3^ box with 0.15 nM NaCl. The system was first energy minimized, with a threshold of max force < 1000 N. Subsequently, NVT (constant number of particles, constant volume, constant temperature) ensemble equilibration and NPT (constant number of particles, constant pressure, constant temperature) ensemble equilibration were performed on the system for 1 ns with a timestep of 1.5 fs respectively. Lastly, a production run was performed on the system. The periodic boundary conditions were applied in all three dimensions. Verlet particle based approach was adopted for neighbor searching with a cutoff distance of 1.2 nm and update frequency of 20 timesteps. The potential-switch method was applied for the short-range Lennard-Jones (LJ) 12-6 interactions from 1 nm to 1.2 nm. The short-range electrostatic interactions were calculated up to 1.2 nm, and the long-range electrostatic interactions were calculated employing the Particle Mesh Ewald algorithm. A time step of 2.5 fs was employed by constraining all the H bonds using the LINCS algorithm. The temperatures were separately coupled for proteins (Fel d 1 and AVA-1-1-C3) and water & ions using the Nosé-Hoover algorithm with the reference temperature 298 K and characteristic time 1 ps.

The isotropic Parrinello-Rahman barostat was used with the reference pressure of 1 bar, the characteristic time of 4 ps, and the compressibility of 4.5 × 10^−5^ bar^-1^. Simulated annealing was applied to the system using a cycling temperature scheme composed of 50 ns windows at constant temperature and 25 ns windows for temperature ramping. The temperature was alternated between a lower bound of 298 K and an upper bound of 350 K. Specifically, the annealing schedule consisted of: Window 1 (0–50 ns) at 298 K; Window 2 (50–75 ns) with a linear ramp from 298 K to 350 K; Window 3 (75–125 ns) at 350 K; and Window 4 (125–150 ns) with a ramp down from 350 K to 298 K. This cycle was repeated for a total simulation time of 1600 ns. Interaction energies were computed using the *gmx energy* module within the GROMACS 2023.5 package. Binding interface RMSDs were calculated using *gmx rms*, with interface residues defined as those within 5 Å of the opposing protein at the binding interface. MD simulations were performed on the AVA-1-1-C3 molecule only to validate the fold structure. The simulations follow the same procedure described above. RMSD was calculated to verify the stability of the structure.

#### Potential of Mean Force Study of binding free energy

After obtaining a stable binding configuration of the Fel d 1–AVA-1-1-C3 complex, umbrella sampling molecular dynamics (MD) simulations were performed to calculate binding free energies. Three systems were investigated: (1) the Fel d 1 tetramer, (2) the Fel d 1 dimer complexed with a single AVA-1-1-C3 molecule, and (3) the Fel d 1 tetramer complexed with two AVA-1-1-C3 molecules. All systems were solvated in 0.15 M NaCl and placed in simulation boxes of appropriate dimensions to ensure convergence of the free energy calculations: 13 × 13 × 20 nm^³^ for the Fel d 1-only system, 13 × 13 × 20 nm^³^ for the Fel d 1 dimer–AVA-1-1-C3 complex, and 20 × 20 × 25 nm^³^ for the Fel d 1 tetramer–AVA-1-1-C3 complex. Each system underwent energy minimization, NVT equilibration, and NPT equilibration using the same parameters as described in the Unbiased MD Simulations section. In the production phase, an external pulling force was applied to selected components: one Fel d 1 dimer in the Fel d 1-only system, the AVA-1-1-C3 molecule in the Fel d 1 dimer–AVA-1-1-C3 complex, and the AVA-1-1-C3–Fel d 1 dimer subcomplex in the Fel d 1 tetramer–AVA-1-1-C3 system. The opposing components were positionally restrained. A simple distance pulling geometry was used, with a pull rate of 0.01 nm/ps and a force constant of 1000 kJ·mol^−¹^·nm^−_2_^. Other simulation parameters remained consistent with those in the unbiased MD simulations. The pulling simulation was run for 1 ns, from which multiple configurations were extracted at 0.2 nm intervals. Each configuration was subjected to a short 100 ps NPT equilibration, followed by a 10 ns production run without any applied pulling force. Free energy profiles were computed using the potential of mean force (PMF), as implemented in gmx wham. Sampling histograms were generated to confirm adequate coverage across the reaction coordinate.

### Size exclusion chromatography

The system of Thermo Vanquish Flex was used to perform size exclusion chromatography. The mobile phase consisted of 150 mM Na_2_HPO_4_, adjusted to pH 7.0. Prior to column installation, the system was equilibrated with the mobile phase. The column (Zenix-C SEC-80, 7.8×300 mm, 3 μm 80 Å) was then connected and further equilibrated by flushing with the mobile phase at a flow rate of 0.7 mL/min until a stable baseline was achieved. All samples were filtered through 0.22 μm membranes and prepared at final concentrations exceeding 0.5 mg/mL. For each run, 30–50 μg of protein was injected at a flow rate of 0.7 mL/min, and UV absorbance at 280 nm was monitored. Following sample analysis, the column was flushed and stored in 50 mM Na_2_PO_4_ containing 0.02% NaN_₃_. To characterize the tetramer disruption of AVA-1-1-C3, the AVA protein or PBS (pH 7.0) and recombinant Fel d 1 was preincubated under 37 ℃ for 4 h under 0.5 mg/mL for both proteins. After preincubation, 30 μL of each mixture was injected for further analysis. The area under the curve of each peak was calculated using python to quantify the dimer and tetramer percentages.

### Der p 1 activity assay

Der p 1 enzyme was pre-activated with 5mM cysteine (Sigma Chemical Co.) to regenerate its thiol group, which becomes oxidized during purification. The catalytic activity of Der p 1 was measured in a continuous rate (kinetic) assay using the fluorogenic peptide substrate N-tert-butoxy-carbonyl (Boc)-Gln-Ala-Arg–7-amino-4-methyl-coumarin (AMC), based on a previous study7,47. The experiment was conducted in 50 mM sodium phosphate buffer, pH 7.0, containing 2.5 mM EDTA and 2.5 mM dithiothreitol (DTT) at 37°C in a total volume of 1 ml. Hydrolysis of AMC substrates was monitored using a Hitachi F-2000 fluorescence with λex = 380 nm and λem = 460 nm. Relative enzymatic activity was then calculated using the following formula:

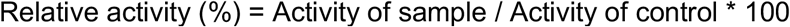

The control (Ctrl) group consisted of reactions containing only the protein and substrate without any inhibitors or modulators. Anti-Der p 1 VHH candidates were prepared at an initial concentration of 0.6 mg/mL in 50 mM phosphate buffer (PB) and subjected to 3-fold serial dilutions in a dilution plate. In a white 96-well assay plate, 10 μL of target antigen (10 μg/mL) was added to each well, followed by the sequential addition of 10 μL of serially diluted antibodies and 10 μL of 10 mM cysteine. The reaction mixtures were incubated at 37 °C for 40 minutes. Subsequently, 80 μL of a 1:1 mixture of substrate solution (50 mM PB, pH 7.0, 10 mM Boc-Gln-Ala-Arg-AMC and 2.5 mM EDTA) and DTT solution (50 mM PB, pH 7.0, 2.5 mM DTT) was added to each well. The plates were immediately transferred to a microplate reader, and fluorescence (λex = 380 nm and λem = 460 nm.) was measured every 10 seconds for 10 minutes. Fluorescence intensity was normalized and used to calculate the percentage inhibition for each VHH candidate.

### In vitro cytotoxicity analysis

Primary human cells were cultured based on the manufacturers-recommended culture media and methods. 1000-2000 cells were plated into 384-well plates and allowed to attach overnight. Varying amounts of AVA or bovine serum albumin (BSA) were added to the cultures for 24-48 hours. CellTiter-Glo® 2.0 was used to measure cell viability using a plate reader. Normalized cell viability was calculated by dividing the luminescence signal by vehicle (PBS) treated cells.

### Thermostability analysis

Thermostability was determined by nano differential scanning fluorimetry (nanoDSF). Tm and Tagg were detected and determined by Prometheus NT.48. The protein samples were diluted to 1 mg/mL into assay buffer and were first centrifuged at 12,000 × g for 10 minutes at 4 °C. After centrifugation, samples were loaded into capillaries. The capillaries were placed sequentially into the corresponding slots, ensuring that each was filled without air bubbles. All samples were heated up from 20 ℃ to 95 ℃ at a constant rate of 1 ℃/min. The ratio of intrinsic fluorescence (350 nm/330 nm) was measured to determine Tm, and Tagg. The experiments were performed in duplicate.

### Statistical analysis

Statistical analyses were performed using Prism v10 (GraphPad) and Python 3.12. Data are reported as mean ± standard deviation (s.d.), unless stated otherwise. All experiments included at least technical duplicates, unless indicated otherwise; biological replicates were performed where feasible. Equality of variances was determined using Levene’s test. For comparisons between groups with equal variances, appropriate parametric tests (e.g., two-tailed Student’s t-test or two-way ANOVA) were applied. A p-value of <0.05 was considered statistically significant.

## Data availability

Data supporting the findings of the study are available within the main manuscript or Supplementary Information. All outstanding data requests may be made to the corresponding author.

## Code availability

All computational analyses and simulation data have been made available at: https://github.com/PacaYang/WhiskerBlockForFeld1.

## Acknowledgements

We would like to thank the team at Pacagen (pacagen.com) for careful review of the manuscript and for important scientific discussions. D.K.Y.Z., Y.L., Z.Q., G.Z., N.A., and E.M.Z. would like to thank the incredible investors in Pacagen for their continued support. The authors would like to thank the many pet parents who advocated for solutions that can foster better living with pets. We are grateful for their commitment to responsible pet adoption and care, which continues to motivate our work and drive the pursuit of new approaches to mitigate allergens in the household.

## Author Information

### Authors and affiliations

Pacagen, Inc, USA

David KY Zhang, Yang Li, Zihan Qu, Gary Zhao, Nathan Andreissen, Evan M Zhao iCareab Biotechnology Co. Ltd., China

Qing Yue, Qiang He

### Contributions

D.K.Y.Z., Q.Y., Q.H., Y.L., and E.M.Z. conceived and designed the experiments. D.K.Y.Z., Q.Y., Q.H., Y.L. and G.Z. performed the experiments. D.K.Y.Z., Q.Y., Q.H., Y.L., Z.Q., N.A., and E.M.Z. analyzed the data. D.K.Y.Z., Y.L., Z.Q., N.A., and E.M.Z. wrote the manuscript. All authors discussed the results and provided comments on the manuscript. The principal investigator is E.M.Z.

## Supporting information

**Supplementary Fig. 1.**
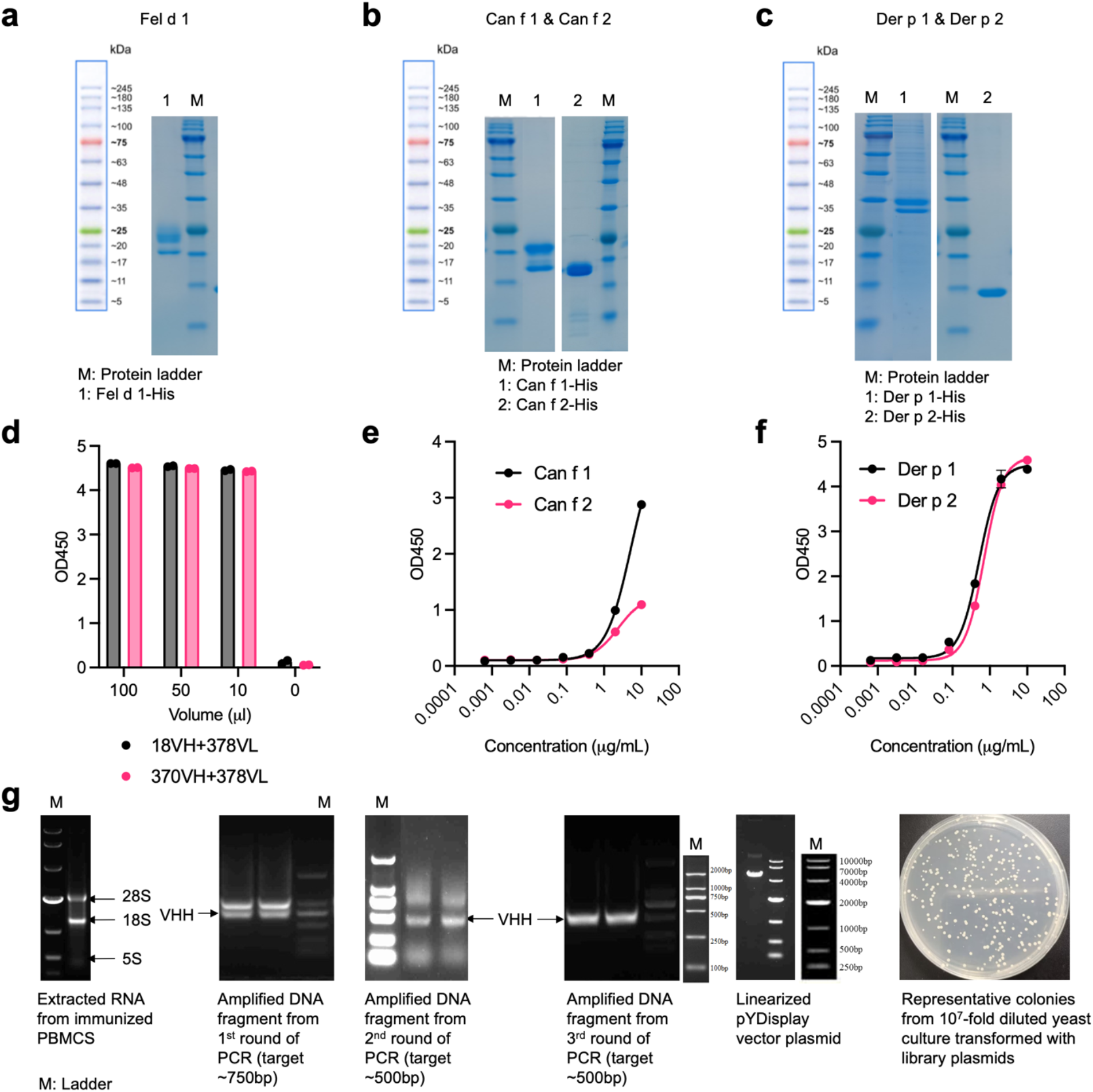
Characterizations of antigens, benchmark antibodies, and constructed libraries. (a-c) SDS-PAGE gels confirmed a >85% purity of recombinant Fel d 1 (a), Can f 1/2 (b), and Der p 1/2 (c). (d-f) Binding activity of purified recombinant antigen proteins and benchmark antibodies was confirmed by ELISA. (d) 18VH+378VL and 370VH+378VL strongly bind to the recombinant Fel d 1. (e) Immunized alpaca serum binds to the recombinant Can f 1 and Can f 2, respectively. (f) Purified 10B9 binds to the recombinant Der p 1, and purified 6OY4 binds to the recombinant Der p 2. (g) Representative construction and diversity determination of the single domain antibody yeast display library. VHH DNA fragments from immunized PBMC were acquired by RNA reverse transcription and PCR amplification. The library diversity was determined to be 3.08×10^9^ by plating the 10^7^-fold diluted transformed yeast. (shown for Fel d 1).

**Supplementary Fig. 2.**
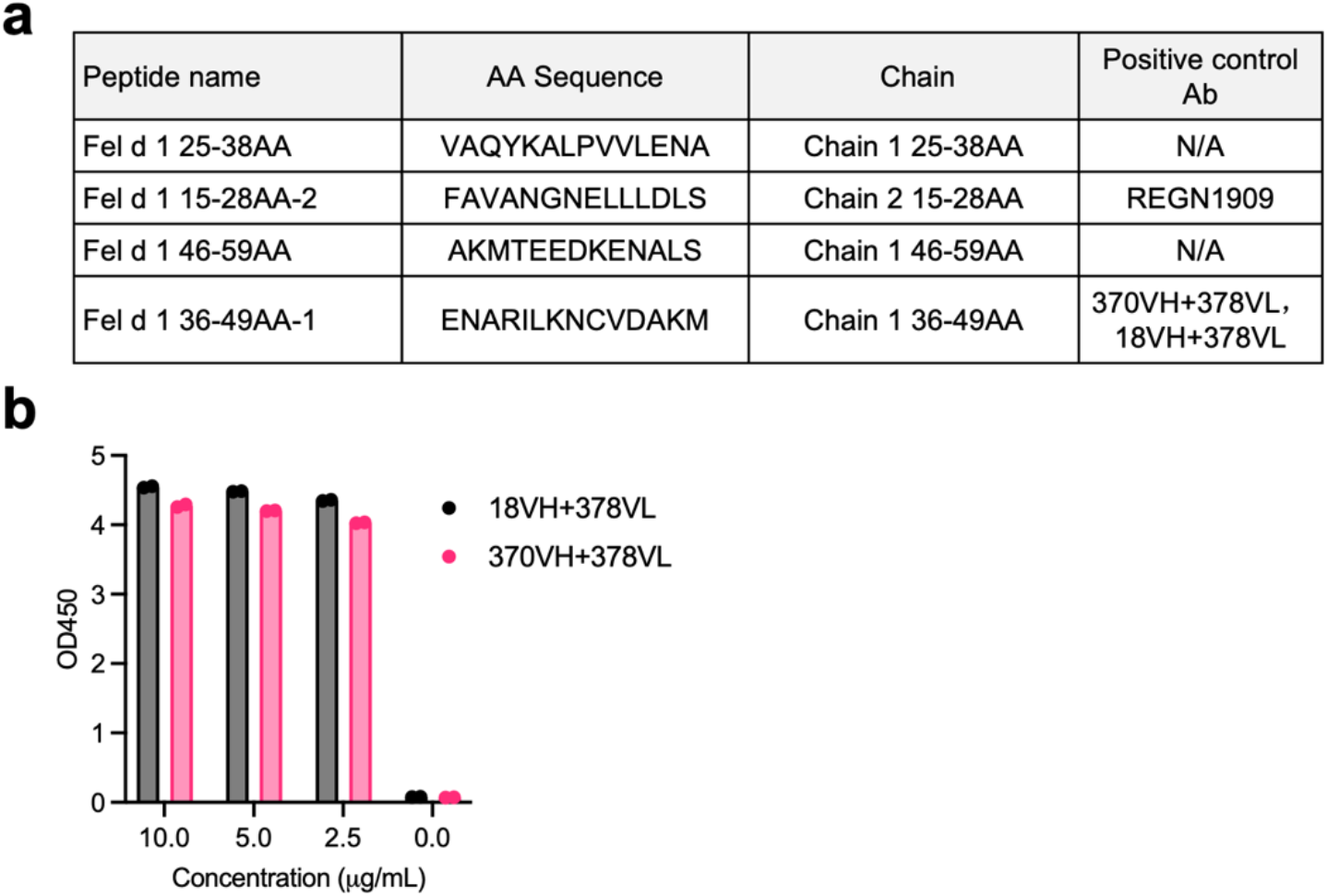
Epitope mapping and diversity characterization of Fel d 1 VHH candidates. (a) the sequences of 4 reported epitopes on Fel d 1 the Fel d 1 protein and corresponding targeting antibodies. (b) Benchmark antibodies 18VH+378VL and 370VH+378VL strongly bind to the epitope Fel d 1 36-49AA-1.

**Supplementary Fig. 3.**
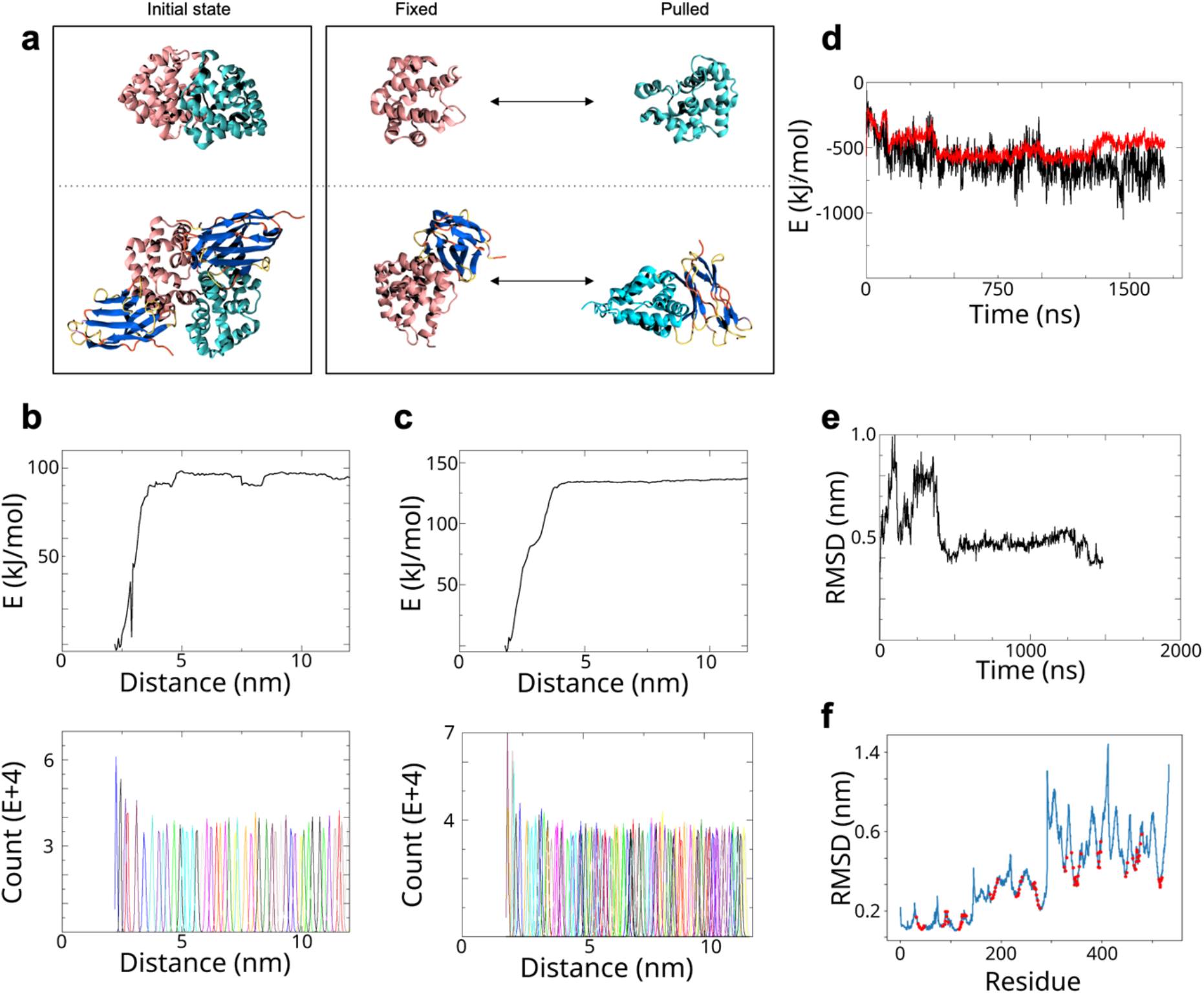
Supporting information for Fel d 1 - AVA computational simulations. (a) Schematic representation of the umbrella sampling molecular dynamics (MD) simulations. Top panel: Fel d 1 tetramer alone; bottom panel: Fel d 1 tetramer in complex with AVA-1-1-C3. (b, c) Results of umbrella sampling MD simulations for the Fel d 1 tetramer without (b) and with (c) AVA-1-1-C3. Top panels show the free energy profiles; bottom panels show the corresponding sampling histograms. (d) Total potential energy of the Fel d 1 tetramer in complex with AVA-1-1-C3. Red: Lennard-Jones interactions; black: Coulombic interactions. (e) Root-mean-square deviation (RMSD) of the binding interface between Fel d 1 and AVA-1-1-C3. (f) Root-mean-square fluctuation (RMSF) of the Fel d 1–AVA-1-1-C3 complex. Highlighted residues indicate the binding interface.

